# Were ancestral proteins less specific?

**DOI:** 10.1101/2020.05.27.120261

**Authors:** Lucas C. Wheeler, Michael J. Harms

## Abstract

Some have hypothesized that ancestral proteins were, on average, less specific than their descendants. If true, this would provide a universal axis along which to organize protein evolution and suggests that reconstructed ancestral proteins may be uniquely powerful tools for protein engineering. Ancestral sequence reconstruction studies are one line of evidence used to support this hypothesis. Previously, we performed such a study, investigating the evolution of peptide binding specificity for the paralogs S100A5 and S100A6. The modern proteins appeared more specific than their last common ancestor (ancA5/A6), as each paralog bound a subset of the peptides bound by ancA5/A6. In the current study, we revisit this transition, using quantitative phage display to measure the interactions of 19,194 random peptides with human S100A5, S100A6, and ancA5/A6. This unbiased screen reveals a different picture. While S100A5 and S100A6 do indeed bind to a subset of the peptides recognized by ancA5/A6, they also acquired new peptide partners outside of the set recognized by ancA5/A6. Our previous work showed that ancA5/A6 had lower specificity than its descendants when measured against biological targets; our new work shows that ancA5/A6 has similar specificity to the modern proteins when measured against a random set of peptide targets. This demonstrates that altered biological specificity does not necessarily indicate altered intrinsic specificity, and sounds a cautionary note for using ancestral reconstruction studies with biological targets as a means to infer global evolutionary trends in specificity.

## Introduction

Changes in protein specificity are essential for evolution (Alhindi *et al.*, 2017; Carroll *et al.*, 2008; Clifton & Jackson, 2016; Kaltenbach & Tokuriki, 2014; Kanzaki *et al.*, 2012; Khersonsky & Tawfik, 2010; Reinke *et al.*, 2013; Soskine & Tawfik, 2010). One intriguing suggestion is that, on average, proteins become more specific over evolutionary time (Copley, 2012; Jensen, 1976; Khersonsky & Tawfik, 2010; Wheeler *et al.*, 2016). If true, this would be a directional “arrow” for protein evolution (Gaucher *et al.*, 2008; Mannige *et al.*, 2012; Risso *et al.*, 2014; Wheeler *et al.*, 2016). Such proposed trends are controversial (Wheeler *et al.*, 2016; Williams *et al.*, 2006), but could ultimately provide fundamental insights into the evolutionary process. For example, increasing specificity might indicate that proteins become less evolvable over time, as they have fewer promiscuous interactions that can be exploited to acquire new functions (Copley, 2012; Khersonsky & Tawfik, 2010). From a practical standpoint, it has also been suggested that less-specific reconstructed ancestors would be powerful starting points for engineering new protein functions (Risso *et al.*, 2013).

There are several reasons that proteins may, on average, evolve towards higher specificity. First, gene duplication followed by subfunctionalization could lead to a partitioning of ancestral binding partners between descendants, and thus increase specificity along each lineage (Alhindi *et al.*, 2017; Clifton & Jackson, 2016; Eick *et al.*, 2012; Hittinger & Carroll, 2007). Second, as metabolic pathways and interaction networks become more complex, proteins must use more sophisticated rules to “parse” the environment: if an ancestral protein had to discriminate between fewer targets than modern proteins, it could be less specific and still achieve the same biological activity (Eick *et al.*, 2012). Finally, on the deepest evolutionary timescales, it has been pointed out that the proteome of the last universal common ancestor was small. As a result, each protein would have been required to perform multiple tasks and hence have lower specificity (Copley, 2012; Jensen, 1976).

Much of the empirical support for the increasing-specificity hypothesis comes from ancestral reconstruction studies (Alhindi *et al.*, 2017; Carroll *et al.*, 2008; Clifton & Jackson, 2016; Devamani *et al.*, 2016; Eick *et al.*, 2012; Ma *et al.*, 2016; Pougach *et al.*, 2014; Rauwerdink *et al.*, 2016; Risso *et al.*, 2013, 2014; Wheeler *et al.*, 2017; Zou *et al.*, 2015). The results from one such study are shown schematically in Fig 1A. We previously studied the evolution of peptide binding specificity in the amniote proteins S100A5 and S100A6. These proteins bind to ≈ 12 amino acid linear peptide regions of target proteins to modulate their activity (Fig 1A) (Bertini *et al.*, 2009; Donato *et al.*, 2013; Leclerc *et al.*, 2009; Lee *et al.*, 2008; Liriano, 2012; Santamaria-Kisiel *et al.*, 2006; S\lomnicki *et al.*, 2009; Streicher *et al.*, 2009). These binding interactions tend to have *K*_*D*_ in the *μM* range. We found that S100A5 and S100A6 orthologs bound to distinct peptides, but that the last common ancestor bound to all of the peptides we tested (Fig 1A) (Wheeler *et al.*, 2017). Other studies, probing other classes of interaction partners, have found similar results: the ancestor interacts with a broader range of partners than extant descendants (Alhindi *et al.*, 2017; Carroll *et al.*, 2008; Clifton & Jackson, 2016; Devamani *et al.*, 2016; Eick *et al.*, 2012; Ma *et al.*, 2016; Pougach *et al.*, 2014; Rauwerdink *et al.*, 2016; Risso *et al.*, 2013, 2014; Wheeler *et al.*, 2017; Zou *et al.*, 2015).

**Fig 1.**
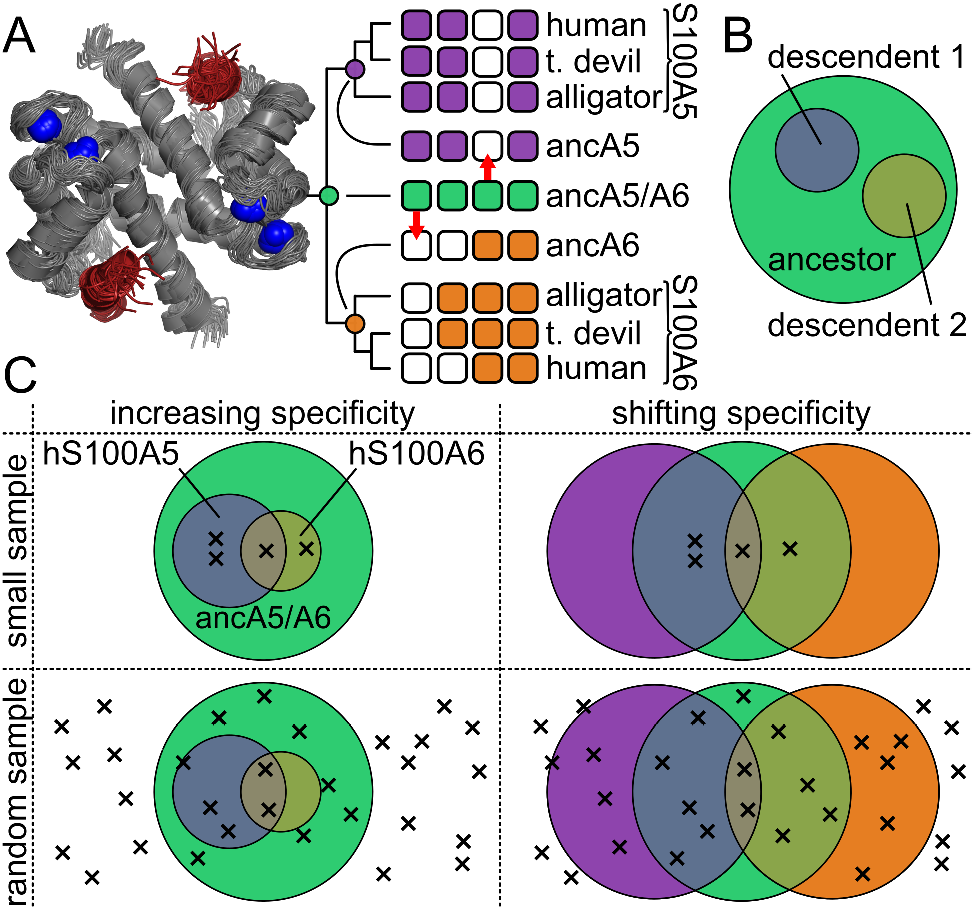
Testing the increased specificity hypothesis requires unbiased sampling of targets. A) Experimentally measured changes in peptide binding specificity for S100A5 and S100A6 (taken from (Wheeler *et al.*, 2017)). Structure: location of peptide (red) binding to a model of S100A5 (gray, PDB: 2KAY). Bound *Ca*^2+^ are shown as blue spheres. Phylogeny: Boxes represent binding of four different peptides (arranged left to right) to nine different proteins (arranged top to bottom). A white box indicates the peptide does not bind that protein; a colored box indicates the peptide binds. Colors denote ancA5/A6 (green), S100A5 (purple), and S100A6 (orange). Red arrows highlight ancestral peptides lost in the modern proteins. B) Venn diagram of the increasing-specificity hypothesis. The large circle is set of targets recognized by the ancestor; the smaller circles are sets of targets represented its descendants. C) Venn diagrams show overlap in peptide binding sets between ancA5/A6, S100A5, and S100A6. Crosses denote experimental observations. Columns show two evolutionary scenarios: increasing specificity (left) versus shifting specificity (right). Rows show to different sampling methods: small sample (top) versus random sampling (bottom). Colors are as in panel B.

But do such experiments truly test the increasing-specificity hypothesis? The hypothesis can be represented as a Venn diagram: the set of targets recognized by the ancestor is larger than the sets of targets recognized by its descendants (Fig 1B). Results such as those in Fig 1A are not, however, sufficient to resolve this Venn diagram. Fig 1C illustrates two radically different Venn diagrams consistent with our experimental observations of peptide binding in Fig 1A. One possibility is increasing specificity (the descendant sets are smaller than the ancestral set). Another possibility is shifting specificity (the descendant sets remain the same size but diverge in their composition). Testing only a small or biased set of binding partners could lead to incorrect conclusions about the evolutionary process. Distinguishing the possibilities shown in Fig 1C requires estimating the populations in each region of the Venn diagram, which can only be done with a much larger, unbiased sample of the set of binding partners.

To test for the evolution of increased specificity, we set out to estimate changes in the total set of peptides between ancA5/A6 and two of its descendants—human S100A5 (hA5) and human S100A6 (hA6). This evolutionary transition is an ideal model to probe this question. We already have a reconstructed ancestral protein that exhibits an apparent gain in specificity over time for both proteins, at least for a small collection of peptides (Wheeler *et al.*, 2017). Further, because they bind to ≈ 12 amino acid peptides, the set of binders is discrete and enumerable (20^12^ = 4 × 10^15^ targets). This contrasts with interactions between proteins and small molecules, for which there is an effectively infinite chemical “space” of possible moieties to sample. We therefore set out to estimate changes in the total sets of partners recognized by these proteins using a high-throughput characterization of peptide binding. We found that the modern proteins bound to a similar number of targets as the ancestor, and that both hA5 and hA6 acquired a large number of new targets since ancA5/A6. Thus, the original observation that a smaller number of targets bound by the ancestor relative to the modern proteins reflects a shifting set of targets—not a shrinking set. This suggests that the evidence for a global trend towards increased specificity from less-specific ancestral states should be revisited.

## Results

### Peptide/protein interactions measured by phage display

Our goal was to measure changes in the total binding sets between human S100A5 (hA5), human S100A6 (hA6), and their last common ancestor (ancA5/A6) (Wheeler *et al.*, 2017). To account for uncertainty in the reconstruction, we also characterized an alternate reconstruction of ancA5/A6 (altAll) that incorporates alternate amino acids at uncertain positions in the reconstruction (Eick *et al.*, 2017). This protein differs at 21 of 86 sites from ancA5/A6, but behaved similary to ancA5/A6 in our previous expereiments (Wheeler *et al.*, 2017).

We first assayed the interaction of tens of thousands of peptides to each protein using phage display. We panned a commercial library of randomized 12-mer peptides expressed as fusions with the M13 phage coat protein. The S100 peptide-binding interface is only exposed upon *Ca*^2+^-binding (Fig 1A); therefore, we performed phage panning experiments in the presence of *Ca*^2+^ and then eluted the bound phage using EDTA (Fig 2A). The population of enriched phage will be a mixture of phage that bind at the site of interest and phage that bind adventitiously (blue and purple phage, Fig 2A). Peptides in this latter category enrich in *Ca*^2+^-dependent manner through avidity or binding at an alternate site (Sidhu *et al.*, 2000; Willats, 2002). To separate these populations, we repeated the panning experiment in the presence of a saturating concentration of competitor peptide known to bind at the site of interest (Fig 2B) (Wheeler *et al.*, 2017). This should lower enrichment of peptides that bind at the site of interest, while allowing any adventitious interactions to remain. By comparing the competitor and non-competitor pools, we can distinguish between actual and adventitious binders.

**Fig 2.**
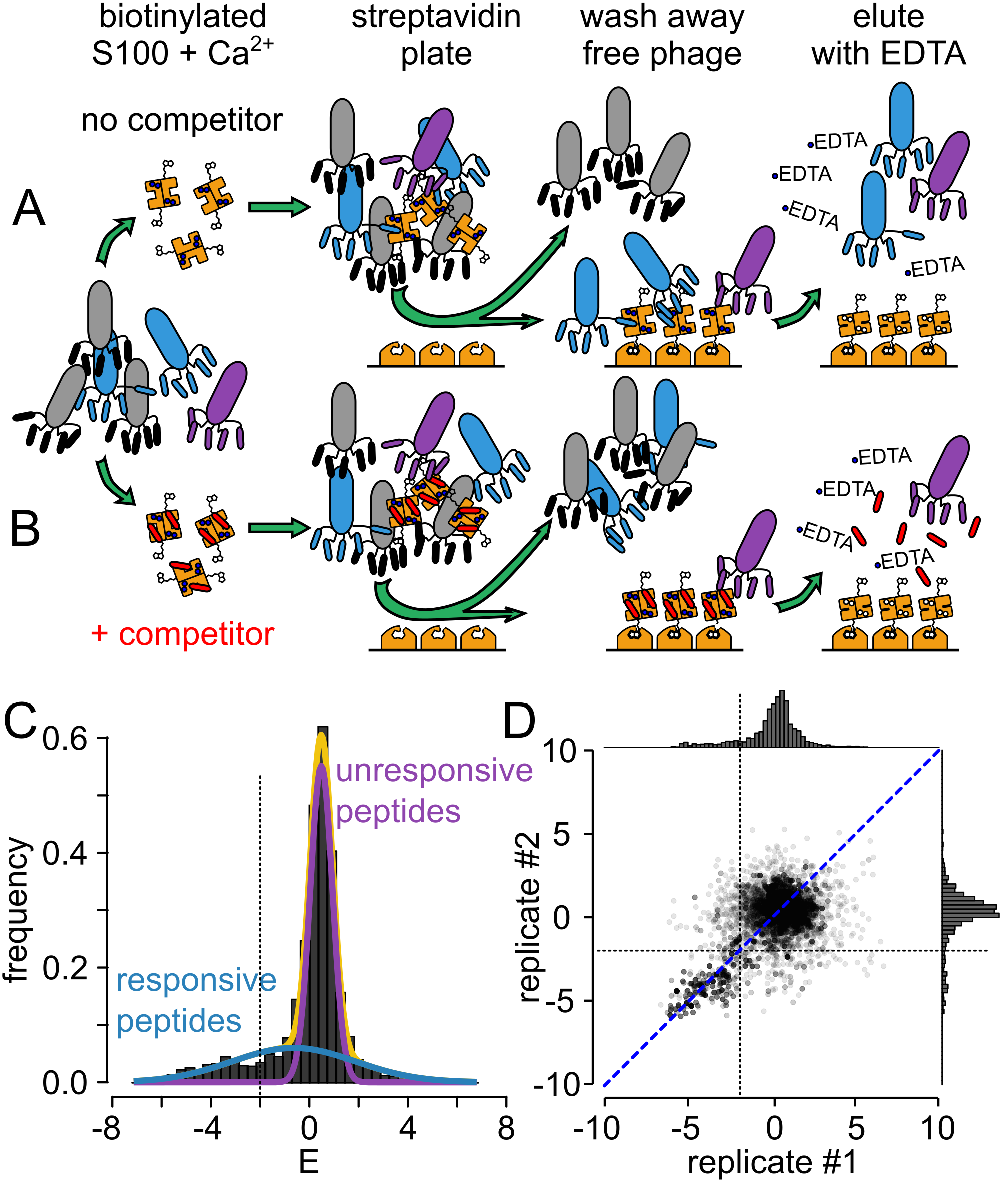
Set of binding peptides can be estimated using phage display. Rows show two different experiments, done in parallel, for each protein. Biotinylated, *Ca*^2+^-loaded, S100 is added to a population of phage either alone (row A) or with saturating competitor peptide added in trans (row B). Phage that bind to the protein (blue or purple) are pulled down using a streptavidin plate. Bound phage are then eluted using EDTA, which disrupts the peptide binding interface. In the absence of competitor (row A), phage bind adventitiously (purple) as well as at the interface of interest (blue). In the presence of competitor (row B), only adventitious binders are present. **C)** Distribution of enrichment values for peptides taken from pooled biological replicates of hA5. The measured distribution (gray) can be fit by the sum of two Gaussian distributions: responsive (blue) and unresponsive (purple), which sum to the total (yellow). The dashed line indicates cutoff for *E* values above which the probability the value arose from the unresponsive distribution is < 0.05. D) Enrichment values from biological replicates are strongly correlated. Axes are enrichment for replicate #1 or replicate #2. Points are individual peptides. Distributions for each replicate are shown on the top and right, respectively. The blue dashed line is the best fit line (orthoganol distance regression), explaining 81% of the variation in the data.

We performed this experiment with and without competitor, in biological duplicate, for all proteins. We found that phage enriched strongly for all proteins relative to a biotin-only control (Fig S1). Further, the addition of competitor binding knocked down enrichment in all samples (Fig S1). After panning, we sequenced the resulting phage pools, as well as the input library, using Illumina sequencing. We applied strict quality control, discarding any peptide that exhibited less than six counts (see methods, Fig S2). After quality control, we had a total of 265 million reads spread over 17 samples (Table S1).

We estimated changes in the frequencies of peptides between samples with and without competitor peptide. For each peptide *i*, we determined *E*_*i*_ = −*ln*(*β*_*i*_/*α*_*i*_), where *β*_*i*_ and *α*_*i*_ are the frequencies of the peptide in the non-competitor and competitor samples, respectively. Defined this way, a more negative value of *E* corresponds to a larger decrease in peptide frequency upon addition of a saturating amount of competitor peptide. More directly: the more negative *E* is for a peptide/protein pair, the better the protein is at selectively enriching the peptide through an interaction at the canonical S100 peptide binding interface. We estimated *E* for ≈ 40, 000 different peptides for each protein (see methods, Fig S3).

We found that the distribution of *E* for each protein could be described using two Gaussian distributions, apparently reflecting two underlying processes (Fig 2C, Fig S4). The dominant peak, centered about *E* = 0, consists of “unresponsive” peptides whose frequencies change little in response to competitor peptide. A second, broader, distribution describes “responsive” peptides whose frequencies change with the addition of competitor. For all proteins, the responsive distribution was shifted towards negative values, meaning the addition of competitor tends to knock off previously bound peptides. Using the mean and standard deviations of the responsive and unresponsive distributions, we could estimate the posterior probability that a peptide with a given value of *E* arose from the responsive rather than unresponsive distribution, and thus reflects specific enrichment of a peptide. We found that *p* = 0.05 occurred around *E* = −1.5 for all proteins (Fig 2C, Fig S4, Fig S5). We therefore interpret peptides exhibiting *E* ≤ −1.5 interacting with the protein at a site that is disrupted by addition of competitor peptide. Values of *E* > −1.5 are exhibited by unreponsive peptides were thus not considered further.

There was no systematic difference between estimates of *E* between biological replicates. We used orthogonal distance regression to compare values of *E* for peptides seen in both biological replicates. The slopes of these lines ranged from 0.9 to 1.1, with intercepts between −0.09 and 0.11. hA5, for example, had a slope 1.06 and an intercept of −0.05 (Fig 2D). There are two distinct regions in these correlation plots, corresponding to the unresponsive and responsive peptide distributions. The unresponsive distribution forms a large cloud about zero. In contrast, the responsive peptide distribution extends along the 1:1 line in a correlated fashion. If we focus on values of *E* < −1.5—peptides arising from the “responsive” distribution—the 1:1 axis of variation explains 87.8% of the total variation in the data for hA5. The worst correlation, observed for altAll, was 75.8%. Finally, because we are comparing changes in the interaction sets between hA5, hA6 and the ancestral proteins, we identified the set of 19,194 peptides that were observed for all four proteins and used this set for our downstream analyses.

### Different changes in specificity occurred along the hA5 and hA6 lineages

We were now in a position to determine how specificity changed over time for these proteins. We first measured the total size of the sets of peptides bound to each protein. If the modern proteins are more specific than their ancestral protein, we would predict they would interact with smaller sets of peptides than the ancestor.

Fig 3A shows the cumulative number of peptides with the *E* below the indicated cutoff. We found that ancA5/A6 interacts with ≈ 1, 400 peptides with *E* ≤ −1.5 (green curve, Fig 3A). Its highest enriching peptide has a score near *E* = −5.5. hA6 has a smaller set of peptides than ancA5/A6 for most values of *E* (orange curve, Fig 3A), in line with the increased specificity prediction. For example, ancA5/A6 has ≈ 500 peptides with *E* ≤ −3, while hA6 has only ≈ 300 peptides over the same interval. hA5, in contrast, has a larger interaction set for all values of *E* (purple curve, Fig 3A). Its highest enriching peptide has *E* = −6.5 and it has ≈ 1, 700 peptides with *E* ≤ −1.5. This indicates that hA5 is not binding a simple subset of the peptides bound by the ancestor.

**Fig 3.**
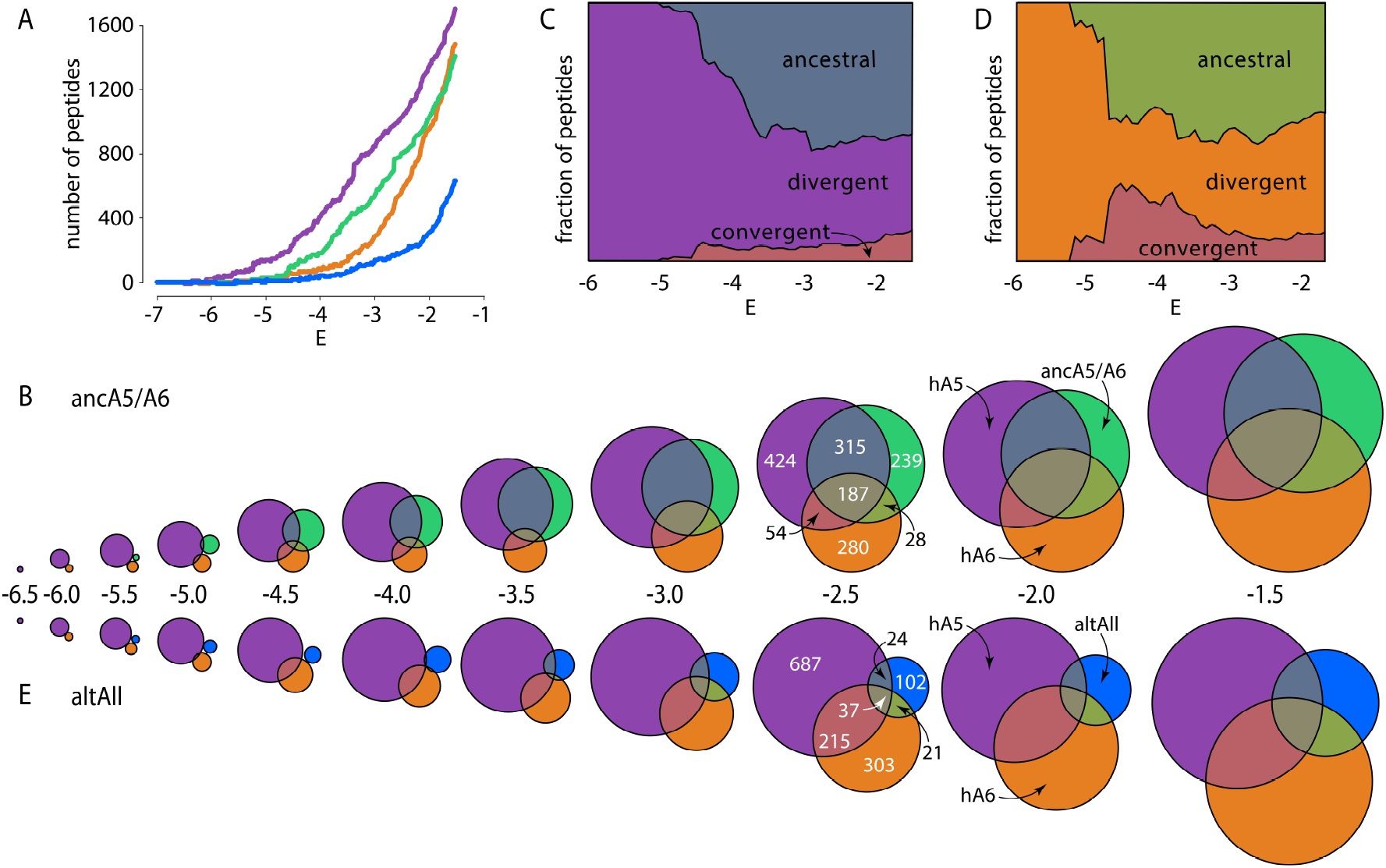
hA5 and hA6 gained new targets since their last common ancestor. A) Cumulative number of peptides observed at or below each enrichment level for hA5 (purple), hA6 (orange), ancA5/A6 (green), and altAll (blue). B) Venn diagrams for overlap between cumulative number peptides observed at or below the indicated enrichment cutoff for hA5 (purple), hA6 (orange) and ancA5A6 (green). The areas are proportional to the number of peptides. The absolute numbers are shown for the *E* ≤ −2.5 Venn diagram for scale. C) Fraction of hA5 targets that overlap with ancA5/A6 (“ancestral”; slate), overlap with neither ancA5/A6 nor hA6 (“divergent”; purple), or overlap with hA6 but not the ancestor (“convergent”; salmon). D) Fraction of hA6 targets that overlap with ancA5/A6 (“ancestral”; green), overlap with neither ancA5/A6 nor hA5 (“divergent”; orange), or overlap with hA5 but not the ancestor (“convergent”; salmon). E) Venn diagrams for overlap between cumulative number peptides observed at or below the indicated enrichment cutoff for hA5 (purple), hA6 (orange) and the altAll ancestor (blue).

To determine if this result was robust to uncertainty in the construction, we also investigated the set of peptides interacting with the alternate version of the reconstructed ancestor, altAll. This protein had a much smaller set of partners than the other three proteins we studied: it only interacts with ≈ 650 peptides with *E* ≤ −1.5. Thus, the result of hA5 gaining targets since the ancestor is robust to phylognetic uncertainty; the result of hA6 becoming more specific is not. If the altAll reconstruction is a better reflection of the evolutionary history of these proteins than ancA5/A6, both hA5 and hA6 appear to have gained new targets over time.

We next probed the overlap between ancA5/A6 and its descendants as a function of *E* (Fig 3B). For the highest enriching peptides—the most negative values of *E*—each protein has a unique set of peptides. At a cutoff of *E* ≤ −5.3, for example, the hA5 set has 76 peptides, the hA6 set has 13 peptides, and the ancA5/A6 set has 15 peptides. None of these peptides overlap between the three protein sets.

At lower stringency *E*, we observed overlap between the peptide sets for the three proteins. By *E* ≤ −2.5, for example, hA5 interacts with 980 peptides. Of these, 424 are unique to hA5 and 502 are shared with ancA5/A6. To determine if this overlap could arise by randomly sampling potential peptide binding partners, we simulated sampling 980 peptides (the size of the hA5 set at this *E* cutoff) and 769 peptides (the size of ancA5/A6 set at this *E* cutoff) from 19,194 unique peptides (the number of peptides in the whole dataset). In these simulations, we observed an average of 39.6 ± 6 shared peptides. The the z-score for observing 502 overlapping peptides is 79.2—giving strong evidence that the overlap is non-random. We did all pairwise comparisons between the sets of hA5, hA6 and ancA5/A6 the Venn diagrams shown in Fig 3B. The worst *p*we found was 2.9 × 10^−7^: the overlapping sets are thus highly significant.

This analysis revealed that hA5, hA6 and ancA5/A6 bind to small, but overlapping, sets of all possible binding peptides (Fig 3B). We next set out to understand the nature of the evolution of these sets. How did hA5 and hA6 change since their last shared common ancestor? A visual inspection of the Venn diagrams reveals that hA5 and hA6 exhibit similar patterns: both have retained many interactions with the ancestral binding set, while also acquring new partners over time. To make better sense of this pattern, we split the interaction targets of hA5 and hA6 into three categories: ancestral (peptides shared with ancA5/A6), convergent (peptides shared between hA5 and hA6, but not the ancestor), and divergent (peptides that are not shared by any others). We then plotted this for hA5 and hA6 as a function of *E* cutoff (Fig 3C,D).

We found that hA5 gave a clear pattern of divergent evolution (Fig 3C). If we look at moderately enriching peptides (−4.0 ≤ *E* ≤ −1.5) we find that ≈ 50% of hA5’s peptides are ancestral, ≈ 45% are divergent, and ≈ 5% are convergent. For the highest enriching peptides (*E* < −4.0), we see the fraction of divergent peptides climbs even higher; for *E* ≤ −5.3, all hA5 peptides were unique and thus—apparently—divergent. One explanation for this observation is a lack of sufficient sampling: maybe the overall fraction of divergent peptides is constant across *E* values, but that the low numbers of peptides with low values of *E* led to a chance over-representation of divergent peptides. To probe for this possibility, we assumed that hA5 had populations like those reflected for moderate enrichment (ancestral: 50%, divgerent: 45% and convergent: 5%). We then walked down *E* and sampled the appropriate numbers of peptides for each *E*. (For example, we sampled a total of 394 hA5 peptides for an *E* cutoff of −4 and only 62 hA5 peptides for an *E* cutoff of −5.5). This allowed us to calculated expected variation in the numbers of peptides in the ancestral, divergent, or convergent categories due to sampling. We found that the observed increase in the relative number of divergent peptides could not be explained by sampling for hA5 (*p* = 5.6 × 10^−16^ for *E* ≥ −5.5; Fig 4A). This indicates that hA5 has acquired an increased number of highly enriching peptides relative to its ancestral state.

**Fig 4.**
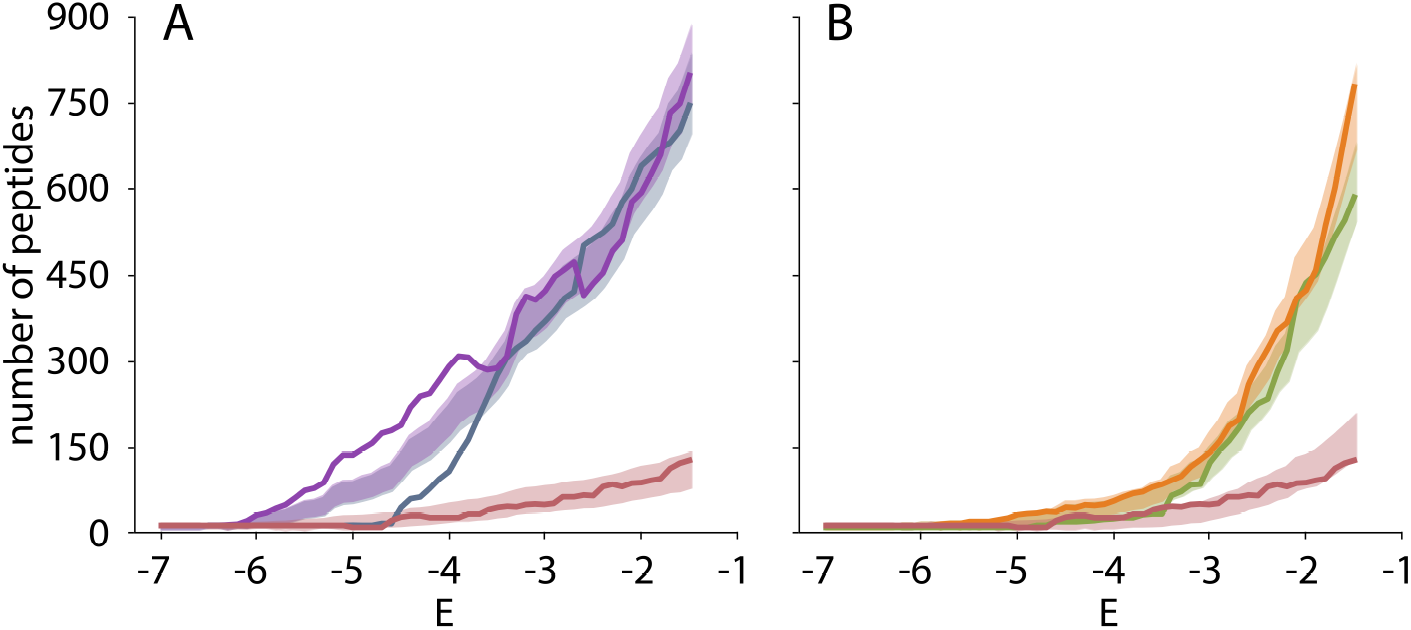
hA5 gained more highly enriched peptides since ancA5/A6. A) Cumulative number of hA5 peptides observed at or below each enrichment level that are ancestral (slate), divergent (purple), or convergent (salmon). The solid lines were observed experimentally (seen in Fig 3C). The shaded regions indicate the 95% confidence intervals for the number of expected peptides if the underlying proportions were ancestral (0.5), divergent (0.45), and convergent (0.05). For *E* < −4.0, the observed number of divergent peptides is elevated above and the observed number of ancestral peptides is depressed below the expectation. B) Equivalent plot for hA6. Curves are ancestral (green), divergent (orange), and convergent (salmon). There is no evidence for elevated numbers of divergent, highly enriched peptides for hA6.

We next turned our attention to hA6. Similar to hA6, it gave a pattern of divergent evolution for moderately enriching peptides (≈ 50% ancestral, ≈ 40% divergent, ≈ 10% convergent) and then climbed to apparently 100% divergent for the most negative values of *E* (Fig 3D). To see if this could be explained by the small numbers of peptides observed at these values of *E*, we repeated the sampling analysis we performed for hA5 for hA6. For hA6, saw no evidence that the relative proportions of divergent, convergent, and ancestral categories changed as a function of enrichment (Fig 4B). Thus, we cannot resolve whether hA6 has acquired a relatively higher proportion of highly enriching targets since ancA5/A6.

Finally, to test the robustness of our conclusions to uncertainty in the reconstruction, we repeated our analysis using the altAll ancestor (Fig 3E). We observed a similar pattern for altAll and ancA5/A6. AltAll interacted with a small set of peptides that partially overlapped with the hA5 and hA6 sets (Fig 3E). AltAll had a much smaller overlap with hA5 and hA6 than ancA5/A6. This small set only exacerbates the trends observed for ancA5/A6. For example, over the interval from −4.0 ≤ *E* ≤ −1.5, an even greater fraction of hA5 peptides are predicted to be divergent (40% for ancA5/A6 vs. 90% for altAll). Likewise, hA6 exhibits a higher proportion of divergent peptides for altAll vs. hA6 (Fig 3E).

Thus, on both lineages, and with two different versions of the reconstructed ancestor, we see that hA5 and hA6 did not gain specificity relative to their ancestral protein. Both lineages maintained interactions with a large number of ancestral protein targets, but also gained a set of new targets. Many of the newly acquired targets were specific to hA5 or hA6, respectively, suggesting a pattern of divergent evolution. The set of partners shifted and grew (hA5) or shifted and slightly shrank (hA6); neither lineage gave a pattern if simple increasing specificity over time.

## Discussion

In this work, we combined ancestral sequence reconstruction with a high-throughput assay to measure evolutionary changes in the specificity of the proteins S100A5 and S100A6. In a previous study of the biological binding partners of the modern proteins, we found that specificity increased since the last common ancestor of the proteins (Fig 1). In the current study, we found the opposite: S100A5, in particular, became less specific over the same evolutionary interval (Fig 3).

These results can be rationalized if we make our definition of specificity more precise. Specificity may be viewed at two levels: *biological* and *instrinsic.* Biological specificity measures the ability of a protein to parse its biological environment. It is determined by both the affinity of a protein for its potential targets and the biological concentrations of the protein and its targets. Such specificity can be tuned by selection, as altering biological interactions can have a profound effect on an organism’s fitness.

Intrinsic specificity, in contrast, measures the affinity of the protein for all possible targets, regardless of whether a given target is encountered by the protein in a biological context. As a whole, intrinsic specificity is invisible to selection: a mutation that alters the ability of a protein to interact with a partner it never encounters will not affect fitness. These latent, promiscuous, interactions can, however, set up future evolutionary change because the preexisting binding interaction can be exploited for new biological functionality (Copley, 2012; Khersonsky & Tawfik, 2010).

A protein with low intrinsic specificity may have a higher latent capacity to form new interactions, potentially making it more “evolvable.” If proteins indeed tend to gain instrinsic specificity over time, one could even argue that they tend to become less evolvable. This would be a striking evolutionary trend (Risso *et al.*, 2013, 2014). Practically, it would also be a strong argument for using reconstructed ancestral proteins as the starting point for protein engineering: a protein with lower instrinsic specificity would have a greater number of latent interactions to exploit and optimize (Khersonsky & Tawfik, 2010; Risso *et al.*, 2018).

Ancestral sequence reconstruction studies have, however, generally probed changes in biological rather than instrinsic specificity (Alhindi *et al.*, 2017; Carroll *et al.*, 2008; Clifton & Jackson, 2016; Devamani *et al.*, 2016; Eick *et al.*, 2012; Ma *et al.*, 2016; Pougach *et al.*, 2014; Rauwerdink *et al.*, 2016; Risso *et al.*, 2013, 2014; Wheeler *et al.*, 2017; Zou *et al.*, 2015). Take our previous study on the evolution of S100A5 and S100A6 (Wheeler *et al.*, 2017). We selected known partners of S100A5 and S100A6, and then asked whether those partners interacted with ancA5/A6. We found that ancA5/A6 bound all of the modern parterns, indicating that S100A5 and S100A6 did, indeed, acquire new biological specificity relative to their ancestor (Fig 1).

The current study, however, reveals that increased biological specificity need not imply an increase in specificity. We found that human S100A5 and S100A6 both acquired many new binding partners over the same interval they acquired new biological specificity. Human S100A5, in particular, binds to *more* targets than its ancestor: it’s intrinsic specificity decreased (Fig 3). This difference between our results for biological and intrinsic specifcity suggests we must carefully define which form of specificity is under discussion when thinking about global evolutionary trends.

Do we expect a trend in global specificity? In our view, the answer is “no.” Because intrinsic specificity is not under selection, the instrinsic specificity of the protein will be determined by chance as the protein meanders through sequence space. Presumably, many more sequences encode low intrinsic specificity than high instrinsic specificity proteins, just as many more sequences encode low stability rather than high stability proteins (Taverna & Goldstein, 2002). As a result, we would expect most evolutionary steps to decrease, rather than increase intrinsic specificity.

Given this, intrinsic specificity would only increase if it was somehow linked to some other feature under selection. One might imagine, for example, that increasing biological specificity necessarily increases intrinsic specificity for certain classes of ligands. It is, however, not obvious that this will generally hold true. A mutation that changes biological specificity alters the chemistry of a protein’s binding interface, excluding some intrinsic partners and adding others. We see no reason to assume the number of partners excluded would be systematically higher than the number added for most classes of mutations and binding sites.

Our work does not rule out a trend of increased intrinsic specificity over deep evolutionary time, but it does caution against interpretating changes in biological specificity as evidence for an overall trend. Testing for a global trend in intrinsic specificity will require studies of unbiased sets of possible interaction partners. It will also necessitates studies of multiple protein families, over deeper evolutionary time scales, and with different classes of binding partners and substrates.

## Materials and Methods

### Molecular cloning, expression and purification in of S100 proteins

Proteins were expressed in a pET28/30 vector containing an N-terminal His tag with a TEV protease cleavage site (Millipore). For each protein, expression was carried out in Rosetta *E.coli* (DE3) pLysS cells. 1.5 L cultures were inoculated at a 1:100 ratio with saturated overnight culture. *E.coli* were grown to high log-phase (*OD*_600_ ≈0.8–1.0) with 250 rpm shaking at 37 °*C*. Cultures were induced by addition of 1 mM IPTG along with 0.2% glucose overnight at 16 °*C*. Cultures were centrifuged and the cell pellets were frozen at 20 °*C* and stored for up to 2 months. Lysis of the cells was carried out via sonication on ice in 25 mM Tris, 100 mM NaCl, 25 mM imidazole, pH 7.4. The initial purification step was performed at 4 °*C* using a 5 mL HiTrap Ni-affinity column (GE Health Science) on an Äkta PrimePlus FPLC (GE Health Science). Proteins were eluted using a 25 mL gradient from 25-500 mM imidazole in a background buffer of 25 mM Tris, 100mM NaCl, pH 7.4. Peak fractions were pooled and incubated overnight at 4 °*C* with ≈1:5 TEV protease (produced in the lab). TEV protease removes the N-terminal His-tag from the protein and leaves a small Ser-Asn sequence N-terminal to the wildtype starting methionine. Next hydrophobic interaction chromatography (HIC) was used to purify the S100s from remaining bacterial proteins and the added TEV protease. Proteins were passed over a 5 mL HiTrap phenyl-sepharose column (GE Health Science). Due to the *Ca*^2+^-dependent exposure of a hydrophobic binding, the S100 proteins proteins adhere to the column only in the presence of *Ca*^2+^. Proteins were pre-saturated with 2mM *Ca*^2+^ before loading on the column and eluted with a 30mL gradient from 0 mM to 5 mM EDTA in 25 mM Tris, 100 mM NaCl, pH 7.4.

Peak fractions were pooled and dialyzed against 4 L of 25 mM Tris, 100 mM NaCl, pH 7.4 buffer overnight at 4 °*C* to remove excess EDTA. The proteins were then passed once more over the 5 mL HiTrap Ni-affinity column (GE Health Science) to remove any uncleaved His-tagged protein. The cleaved protein was collected in the flow-through. Finally, protein purity was examined by SDS-PAGE. If any trace contaminants appeared to be present we performed anion chromatography with a 5 mL HiTrap DEAE column (GE). Proteins were eluted with a 50 mL gradient from 0-500 mM NaCl in 25 mM Tris, pH 7.4 buffer. Pure proteins were dialyzed overnight against 2L of 25 mM TES (or Tris), 100 mM NaCl, pH 7.4, containing 2 g Chelex-100 resin (BioRad) to remove divalent metals. After the final purification step, the purity of proteins products was assessed by SDS PAGE and MALDI-TOF mass spectrometry to be > 95. Final protein products were flash frozen, dropwise, in liquid nitrogen and stored at −80 °*C*. Protein yields were typically on the order of 25 mg/1.5 L of culture.

### Preparation of biotinylated proteins for phage display

A mutant version of hA5 with a single N-terminal Cys residues were generated via sitedirected mutagenesis using the QuikChange lightning system (Agilent). The Cys was introduced in the Ser-Asn tag leftover from TEV protease cleavage as Ser-Asn-Cys. The proteins were expressed and purified as described in the previous section. A small amount of the purified proteins were biotinylated using the EZ-link BMCC-biotin system (ThermoFisher Scientific). ≈1 mg BMCC-biotin was dissolved directly in 100% DMSO to a concentration of 8 mM for labeling. Proteins were exchanged into 25mM phosphate, 100mM NaCl, pH 7.4 using a Nap-25 desalting column (GE Health Science) and degassed for 30 min at 25 °*C* using a vacuum pump (Malvern Instruments). While stirring at room temperature, 8mM BMCC-biotin was added dropwise to a final 10X molar excess. Reaction tubes were sealed with PARAFILM (Bemis) and the maleimide-thiol reactions were allowed to proceed for 1 hour at room temperature with stirring. The reactions were then transferred to 4°*C* and incubated with stirring overnight to allow completion of the reaction. Excess BMCC-biotin was removed from the labeled proteins by exchanging again over a Nap-25 column (GE Health Science), and subsequently a series of 3 concentration-wash steps on a NanoSep 3K spin column (Pall corporation), into the Ca-TeBST loading loading buffer. Complete labeling was confirmed by MALDI-TOF mass spectrometry by observing the ≈540Da shift in the protein peak. Final stocks of labeled proteins were prepared at 10 *μM* by dilution into the loading buffer.

### Phage display

Phage display experiments were performed using the PhD-12 peptide phage display kit (NEB). All steps involving the pipetting of phage-containing samples was done using filter tips (Rainin). We prepared 100 *μL* samples containing phage (5.5 × 10^11^ PFU) and 0.01 *μM* biotin-protein (or biotin alone in the negative control) at room temperature in a background of *Ca*^2+^-TeBST loading buffer (50mM TES, 100mM NaCl, 2mM *CaCl*_2_, 0.01% Tween-20, pH 7.4) to ensure *Ca*^2+^-saturation of the S100 proteins. For the experiments using a peptide competitor, we included the peptide RSHSGFDWRWAMEALTGGSAE at 20 *μM* in the loading buffer. This peptide (named A6cons in the original report), binds all four proteins at the canonical binding site with *K*_*D*_ between 1 and 8 *μM* (Wheeler *et al.*, 2017). Samples were incubated at room temperature for 2hr. Each sample was then applied to one well of a 96-well high-capacity streptavidin plate (previously blocked using PhD-12 kit blocking buffer and washed 6X with 150 *μL* loading buffer). Samples were incubated on the plate with gentle shaking for 20min. 1 *μL* of 10 *mM* biotin (NEB) was then added to each sample on the plate and incubated for an additional five minutes to compete away purely biotin-dependent interactions. Samples were then pulled from the plate carefully by pipetting and discarded. Each well was washed 5X with 200 μ*L* of loading buffer by applying the solution to the well and then immediately pulling off by pipetting. Finally, 100 μ*L* of EDTA-TeBST elution buffer (50mM TES, 100mM NaCl, 5mM EDTA, 0.01% Tween-20, pH 7.4) was applied to each well and the plate was incubated with gentle shaking for 1hr at room temperature to elute. Eluates were pulled from the plate carefully by pipetting and stored at 4°*C*. Eluates were titered to quantify eluted phage as follows. Serial dilutions of the eluates from 1 : 10 − 1 : 10^5^ were prepared in LB medium. These were used to inoculate 200 μ*L* aliquots of mid-log-phase ER2738 *E. coli* (NEB) by adding 10 μ*L* to each. Each 200 μ*L* aliquot was then mixed with 3mL of pre-melted top agar, applied to a LB agar XGAL/IPTG (Rx Biosciences) plate, and allowed to cool. The plates were incubated overnight at 37°*C* to allow formation of plaques. The next morning, blue plaques were counted and used to calculate PFU/mL phage concentration. Enrichment was calculated as a ratio of experimental samples to the biotin-only negative control.

To generate the input phage library, the commercially-produced library was first screened in duplicate against each of the four proteins as described above. Each of these lineages was subsequently amplified in ER2738 *E. coli* (NEB) as follows. 20mL 1:100 dilutions of an ER2738 overnight culture were prepared. Each 20mL culture was inoculated with one entire sample of remaining phage eluate. The cultures were incubated at 37°*C* with shaking for 4.5 hours to allow phage growth. Bacteria were then removed by centrifugation and the top 80% of the culture was removed carefully with a filtered serological pipette and transferred to a fresh tube containing 1/6 volume of PEG/NaCl (20% w/v PEG-8000, 2.5M NaCl). Samples were incubated overnight at 4°*C* to precipitate phage. Precipitated phage were isolated by centrifugation and subsequently purified by an additional PEG/NaCl precipitation on ice for 1hr. These individually amplified pools were then resuspended in 200 μ*L* each of sterile loading buffer and mixed together to form a pre-conditioned library in order to minimize the impact of sampling on the subsequent panning experiment. The pool was diluted 1:1 with 100% glycerol and stored at −20°*C* for use in the final panning experiments.

### Preparation of deep sequencing libraries

Phage genomic ssDNA was isolated from leftover amplified eluates from each round of panning using the M13 spin kit (Qiagen). Products were stored in low TE buffer. These ssDNA were used as the template for 2 replicate PCRs with the Cs1 forward (5’—ACACTGACGA-CATGGTTCTACAGTGGTACCTTTCTATTCTCACTCT—3’) and PhD96seq-Cs2 reverse (5’—TACGGTAGCAGAGACTTGGTCTCCCTCATAGTTAGCGTAACG—3’) primers. Pro ucts were isolated from these PCR products using the GeneJet gel extraction kit (Thermo Scientific) and pooled. The pooled products were then used as templates for a secondary reaction with the barcoded primers. Products were isolated from these final PCRs using the GeneJet gel extraction kit. Concentration of barcoded samples was measured by *A*_260_*/A*_280_ using a 1mm cuvette on an Eppendorf biospectrometer. Multiplexing was done by mixing samples according to mass. The concentration of the multiplexed library was corrected using qPCR with the P5 and P7 Illumina flow-cell primers. The library was then diluted to a final concentration of 10nM and Illumina sequenced on two lanes of a HiSeq 4000 instrument, using the Cs1 F’ as the R1 sequencing primer. The lanes were spiked with 20% PhiX control DNA due to the relatively low diversity of the library.

### Phage display analysis pipeline

We performed quality control on three read features. First, we verified that the sequence had exactly the anticipated length from the start of the phage sequence through the stop codon. Second, we only took sequences in which the invariant phage sequence differed by at most one base from the anticipated sequence. This allows for a single point mutation and or sequencing errors, but not wholesale changes in the sequence. Finally, we took only reads with an average phred score better than 15. The vast majority of the reads that failed our quality control did not have the variable region, representing reversion to phage with a wildtype-like coat protein. This analysis is encoded in the *hops_count.py* script (https://github.com/harmslab/hops), which takes a gzipped fastq file as input and returns the counts for every peptide in the file.

### Identifying the read count cutoff

One critical question is at what point the number of reads correlates with the frequency of a peptide. If we set the cutoff too low, we incorporate noise into downstream analyses. If we set the cutoff too high, we remove valuable observations from our dataset. To identify an appropriate cutoff, we studied the mapping between *c*_*i*_ (the number of reads arising from peptide *i*) and *f*_*i*_ (the actual frequency of peptide *i* in the experiments). Our goal was to find *P*(*f*_*i*_|*c*_*i*_, *N*): the probability peptide *i* is at *f*_*i*_ given we observe it *c*_*i*_ times in *N* counts. Using Bayes theorem, we can write

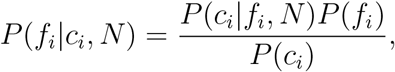

where *N* is the total number of reads. We calculated *P*(*c*_*i*_|*f*_*i*_, *N*) assuming a binomial sampling process: what is the probability of observing exactly *c* counts given *N* independent samples when a population with a peptide frequency *f*_*i*_? This gives the curve seen in Fig S2A. We then estimated 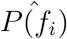 from the distribution of frequencies in the input library, constructing a histogram of apparent peptide frequencies (Fig S2B). Empirically, we found that frequencies followed an exponential distribution over the measurable range of frequencies. Finally, we assumed that all counts have equal prior probabilities, turning *P*(*c*_*i*_) into a scalar that normalizes the integral of *P*(*f*_*i*_|*c*_*i*_, *N*) so it sums to 1.

Using the information from Fig S2A and B, we could then calculate *P*(*f*_*i*_|*c*_*i*_, *N*) for any number of reads in an experiment *N* . Fig S2C shows this calculation for *N* = 2.0 × 10^7^ reads—a typical number of reads from our experimental replicates. This curve is linear above 6 reads. Below this, counts no longer correlates linearly with frequency, as it is possible to obtain 5 reads random sampling from low frequency library members. We therefore used a cutoff of 6 counts for all downstream analyses. In total, 74.0% of reads passed our quality control and read cutoff (Table S1).

### Measuring enrichment values

We next set out to measure changes in the frequency of peptides between the competitor and non-competitor samples. The simplest way to do this would be to identify peptides seen in both experiments, and then measure how their frequencies change between conditions. Unfortunately, these proteins all bind a wide swath of peptide targets and relatively few peptides were shared between conditions. This approach would thus exclude the majority of sequences. For example, only 8,672 of the 112,681 unique peptides observed for hA5 were present in both the competitor and non-competitor, even after pooling biological replicates. Worse, because we are interested in peptides that are lost when competitor peptide is added, ignoring peptides with no counts in the competitor sample means ignoring some of the most informative peptides.

To solve this problem, we clustered similar peptides and measured enrichment for peptide clusters rather than individual peptides. We extracted all peptides that were observed across the competitor and non-competitor samples for a given protein. We then used DBSCAN to cluster those peptides according to sequence similarity, as measured by their their Damerau-Levenshtein distance (Damerau, 1964; Ester *et al.*, 1996). This revealed extensive structure in our data. For example, hA5 yielded 8,645 clusters with more than one peptide, incorporating more than half of the unique peptides (Fig S3A). We chose clustering parameters that led to highly similar peptides within each cluster, as can be seen by the representative sequence logos for three clusters of hA5 (Fig S3B). Sequences that were not placed in clusters were treated as clusters with a size of one.

We then used the enrichment of each cluster to estimate the enrichment of individual peptides. We defined enrichment as:

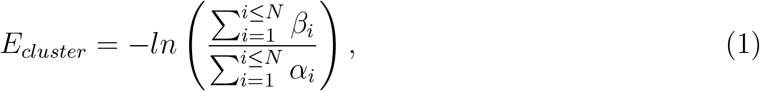

where *N* is the total number of peptides in the cluster, *β*_*i*_ is the frequency of peptide *i* in the competitor sample, and *α*_*i*_ is the frequency of peptide *i* in the non-competitor sample. We then made the approximation that all members of the cluster have the same enrichment:

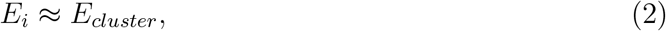

allowing us to estimate the enrichment of all *i* peptides in the cluster (Fig S3C). Peptides lost because of competition for the interface will add zeros to the numerator of Eq. 1, leading to an overall decrease in enrichment. Peptides missed because of finite sampling will add zeros evenly to the competitor and non-competitor samples, leading to no net enrichment.

We tested this cluster-based approximation using the 8,672 peptides of hA5 for which we could directly calculate enrichment (that is, those peptides seen in both the competitor and non-competitor experiments). We calculated the enrichment of each peptide individually and compared these values to those obtained by the cluster method. There is no systematic difference in the values estimated using the two methods, and the linear model explains 98.4% of the variation between the two methods.

We clustered peptides using our own implementation of the DBSCAN algorithm (Ester *et al.*, 1996) using the Damerau-Levensthein distance (Damerau, 1964). The main parameter for DBSCAN clustering is *ε*—the neighborhood cutoff. Clusters are defined as sequences that can be reached through a series of *ε*-step moves. We found that *ε* = 1 gave the best results for our downstream machine learning analysis. Our whole enrichment pipeline—including clustering—can be run given a peptide count file for the non-competitor experiment and a peptide-count file for the competitor experiment using the *hops_enrich.py* script (https://github.com/harmslab/hops).

## Acknowledgements

We’d like to thank members of the Harms lab for critical feedback on the manuscript. This work was supported by NIH R01GM117140 (MJH) and NIH 7T32GM007759-37 (LCW).

---

**Fig S1.**
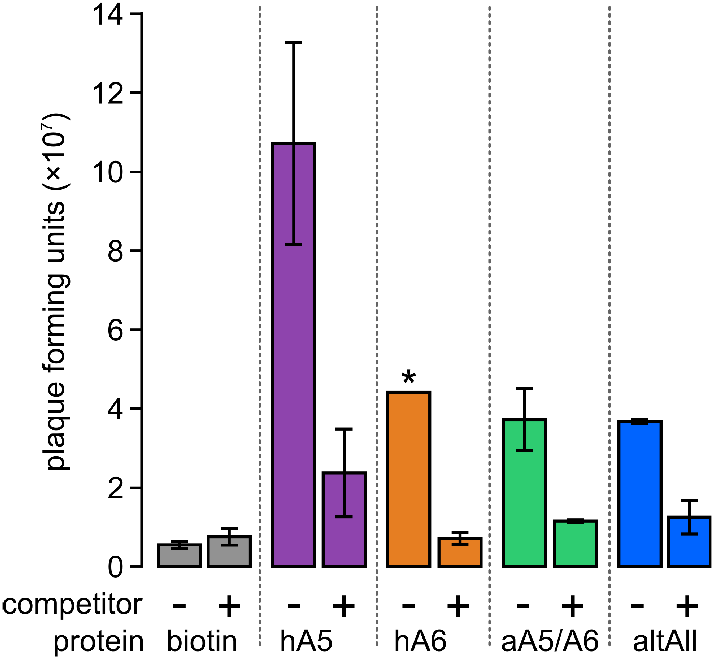
Phage enrichment is reduced in the presence of competitor peptide. Figure shows eluted plaque forming units (PFU) (estimated from phage titer) for two biological replicates of each condition. Enrichment is shown for biotin-only control (gray), hA5 (purple), hA6 (orange), ancA5/A6 (dark green), and altAll (light green) with (+) and without (−) 20 *μM* competitor peptide. Error bars show the standard error for two biological replicates. (*) hA6 without competitor is shown for only one replicate due to failure of the titer for the one replicate.

**Fig S2.**
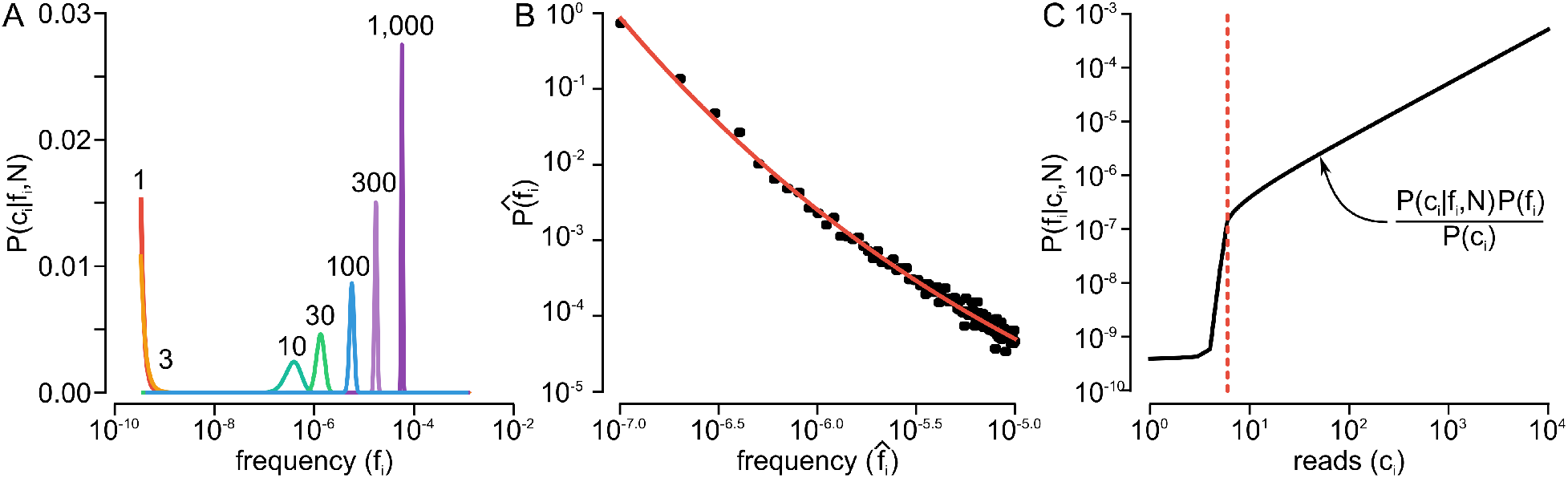
We can identify the number of counts that reliably reports on frequency in a sequenced phage pool. A) Using binomial sampling, we can calculate the probability of observing exactly *c*_*i*_ counts in *N* samples from a pool that has a peptide of actual frequency *f*_*i*_. Figure shows curves for counts ranging from 1 (red) to 1,000 (pink), all using *N* = 2.0×10^7^. B) Panel shows a histogram of frequencies estimated from 3.9×10^7^ reads taken from the input library. The black points are experimental data. The red curve is an exponential distribution fit to that curve. C) Using the sampling from panel A and the fit curve from panel B, we can determine *P*(*f*_*i*_|*c*_*i*_, *N*). The solid curve shows the relationship between the number of reads for peptide *i* (x-axis) against the maximum-likelihood estimate of the frequency (y-axis). The red line highlights the cutoff we used in our experiments.

**Fig S3.**
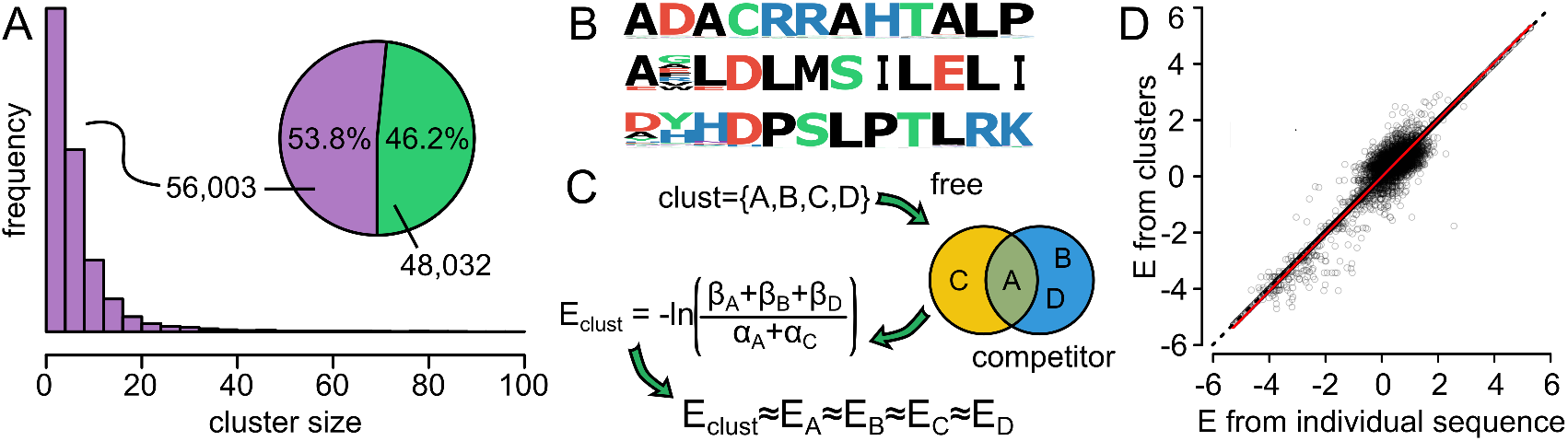
We can estimate how addition of competitor peptide alters the frequencies of peptides. A) Distribution of sizes of peptide clusters from hA5 experiment. Pie chart shows number of peptides placed in clusters (56,003; 53.8%) versus not (48,032; 46.2%). B) Three example clusters taken from the clusters in panel A. The letter height at each position indicates its frequency in the sequences within that cluster. C) Toy example showing how enrichment is calculated for a cluster containing peptides {*A*, *B*, *C*, *D*}. Peptides *A* and *C* were observed in the no competitor sample at frequencies *α*_*A*_ and *α*_*C*_ . Peptides *A*, *B*, and *D* were observed in the competitor sample at frequencies *β*_*A*_, *β*_*B*_ and *β*_*D*_. The enrichment of the cluster is given by *E*_*clust*_ = −*ln*[(*β*_*A*_ + *β*_*B*_ + *β*_*D*_)/(*α*_*A*_ + *α*_*C*_)]. All members of the cluster are then assigned *E* ≈ *E*_*clust*_. D) Comparison of enrichment values for hA5 peptides determined using a direct comparison of frequencies with and without competitor (x-axis) versus the clustering method (y-axis). Each point is an individual peptide. Red line is a least-squares regression line fit to the data. The dashed line is the 1:1 line.

**Fig S4.**
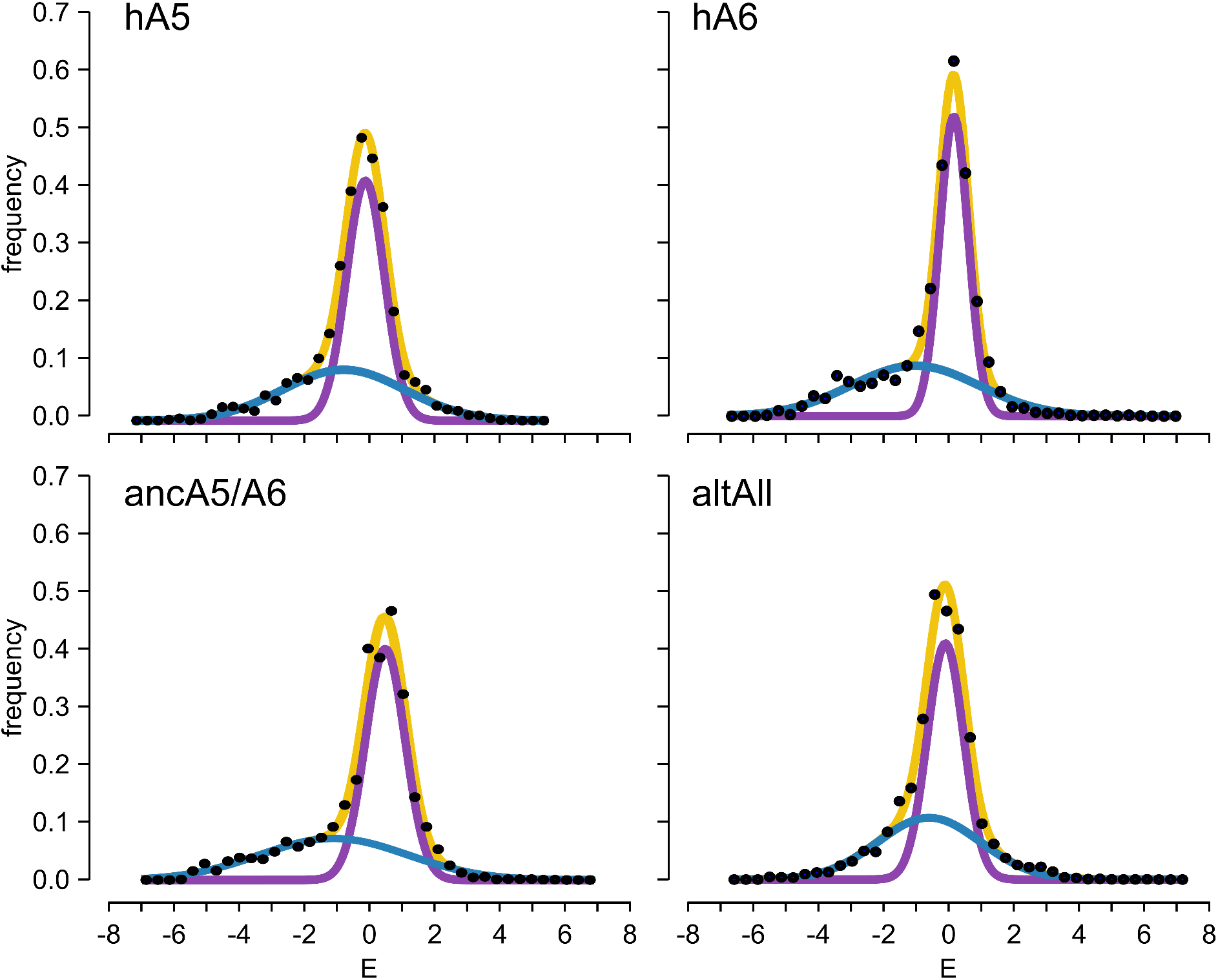
Distributions of *E* for each protein. Panels show distribution of *E* for each protein (pooled bio-replicates). Points are raw histograms. Curves are two Gaussian fit: blue (responsive), purple (unresponsive) and yellow (sum).

**Fig S5.**
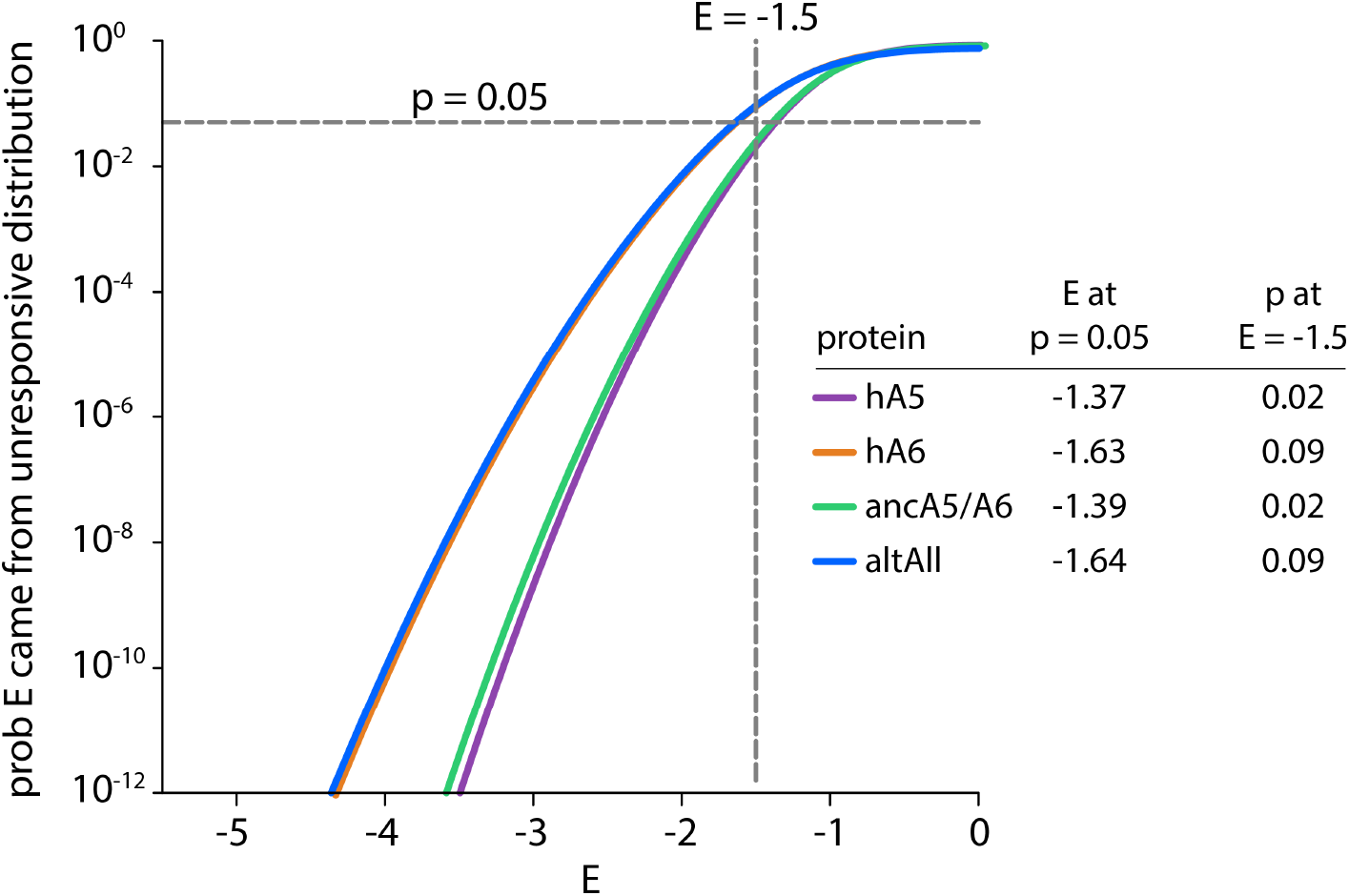
Selection of *E* = −1.5 as a cutoff for peptide enrichment. Curves show the prosterior probability a peptide with the enrichment score shown on the x-axis arose from the “unresponsive” distribution for each protein. These curves were calculated by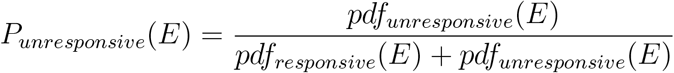where *pdf* is the probability density function in *E* for the normal distributions seen in Fig 2C and S4. The inset table shows the *E* value where the *P*_*unresponsive*_(*E*) is 0.05 and the *P*_*unresponsive*_(*E*) at *E* = −1.5.

**Table S1:** Number of sequencing reads for each sample. Sample, and whether or not competitor was added, are indicated on the right. Columns show biological replicates 1 or 2. “total” columns indicate reads returned by Illumina software pipeline. “good” columns indicate reads that passed our quality control and were used to calculate enrichment values.

| sample | competitor | rep1 |  | rep2 |  |
| --- | --- | --- | --- | --- | --- |
|  |  | total | good | total | good |
| hA5 | - | 24,794,016 | 19,695,958 | 29,085,203 | 16,773,567 |
| hA5 | + | 15,053,706 | 11,523,991 | 17,631,137 | 13,612,463 |
| hA6 | - | 22,728,393 | 17,722,779 | 7,769,003 | 5,972,295 |
| hA6 | + | 13,953,466 | 11,004,701 | 23,026,469 | 18,128,759 |
| raw library |  | 39,700,991 | 32,190,368 | — | — |

